# A Transferable and Robust Computational Framework for Class A GPCR Activation Free Energies

**DOI:** 10.64898/2025.12.05.692536

**Authors:** Simone Aureli, Nicola Piasentin, Thorben Fröhlking, Valerio Rizzi, Francesco Luigi Gervasio

## Abstract

The activation of G-protein coupled receptors is involved in many bio-medically important cellular pathways. However, capturing it with molecular simulations is far from trivial as it requires capturing both local and global motions. We recently achieved this goal in a specific receptor (the *β*1-adrenergic receptor, or ADRB1) by combining a multiple replica enhanced sampling approach with tailored collective variables. While that approach can be applied to other receptors, it would require a tedious and error-prone choice and refinement of the collective variables, and in particular of the main path-like variable. Herein, we introduce an effective and streamlined evolved strategy for defining the CVs that reduces user intervention while still achieving a robust free energy convergence. We apply it to two apo-GPCRs of pharmacological relevance, ADRB1 and the *µ*-opioid receptor. In the first case we show that the reconstructed free energies agree with those obtained with the previous tailored approach, while for the *µ*-opioid receptor activation we gain novel biological insights. The proposed method can be easily applied to other class A GPCRs, paving the avenue to the systematic elucidation of the activation mechanisms of many crucial drug targets.

**TOC Graphic:** 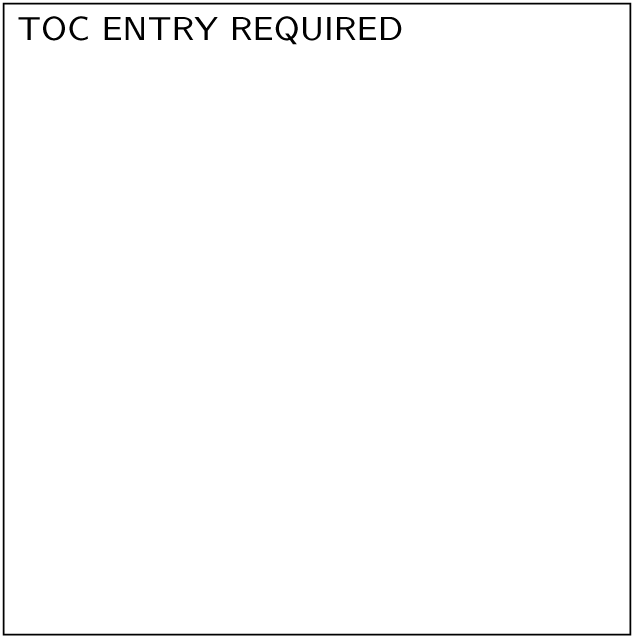

## 1 Introduction

G protein–coupled receptors (GPCRs) are one of the most important classes of membrane proteins, responsible for transducing extracellular signals into intracellular responses and serving as targets for a large fraction of approved drugs^1–3^. Their function is tightly linked to conformational changes that span multiple scales, from local rearrangements of side chains and hydration sites to large-scale reorganizations of the transmembrane helices that define the transition between inactive and active states^4,5^.

Several simulation strategies have been applied to investigate GPCR activation, ranging from unbiased molecular dynamics (MD) simulations to various enhanced sampling methods^6–17^. While unbiased MD simulations have revealed interesting details ^6–8,18^, receptors’ transitions occur on timescales that remain difficult to access with unbiased MD simulations, a limitation that is commonly addressed by enhanced sampling strategies^12,14–16,19,20^. In particular, collective variable (CV)-based approaches aim to accelerate the reaction coordinates associated with the key processes under investigation^21–24^.

Among them, our carefully designed OneOPES framework enables simultaneous acceleration of multiple CVs, offering an efficient way to combine descriptors of both microscopic and mesoscopic motions within a single simulation^25^. Recently, this strategy has been successfully applied to the activation of *β*1-adrenergic receptor (ADRB1), where specifically tailored CVs captured both local changes such as side-chain rearrangements and hydration dynamics, as well as the large-scale conformational reorganization of the receptor backbone^17^.

Within the OneOPES approach, the path collective variable (PATH CV, hereafter) has proven to be a natural and effective choice to represent GPCRs’ substantial backbone rearrangements. First introduced nearly twenty years ago^26^, the original PATH CV requires the definition of a sequence of representative intermediate structures that bridge two target states, ideally arranged at regular intervals of RMSD. In practice, this setup can be labor-intensive, i.e., multiple manually curated structural hypotheses must be generated, tested, and refined in a trial-and-error process until a stable and physically meaningful path is obtained. This limitation reduces the accessibility and transferability of the method, making it challenging to apply it across multiple receptors or conformational transitions systematically.

Here, we present a robust and transferable strategy that streamlines the construction of PATH CVs, substantially reducing the manual intervention required while preserving the original accuracy. It is critical to emphasize that this approach only requires knowledge of the two target structures, eliminating the necessity to manually define and optimize a sequence of intermediate structures. We validate the approach on two pharmacologically relevant GPCRs in their apo forms, i.e., the *β*1-adrenergic receptor (ADRB1) and the *µ*-opioid receptor (MOR). For both systems, the refined PATH CV reproduces the free-energy landscapes obtained with the original tailored PATH CV approach, but with shorter simulation times needed to achieve convergence and substantially reduced setup effort. This evolution thus improves both the efficiency and the transferability of the OneOPES sampling protocols for GPCR activation, and more broadly, for other biomolecular systems undergoing large-scale conformational transitions.

## 2 Methods

### Definition of the Euclidean PATH CV

The original RMSD-based PATH CV formulation^26^, from now on referred to as RPATH, requires the definition of a sequence of representative intermediate conformations linking inactive and active states. This provides a *progress* variable *s* and a *deviation* variable *z* defined as follows:

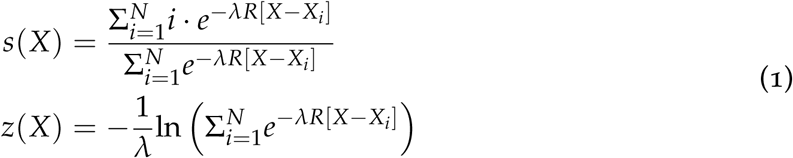

In this formulation, the path is described by a set of *N* reference structures or milestones *X_i_*, and the variables *s* and *z* are expressed as functions of the RMSD distances *R*[*X − X_i_*] between the current configuration *X* and each of these high-dimensional reference frames *X_i_*. The parameter *λ* determines the smoothness of the projection on the reference frames, with high *λ* values rigidly tying *s* to the closest single reference frame and producing a typical stepwise behavior in the dynamics, while low *λ* values spreading out *s* over a number of reference frames. In enhanced sampling, the sweet spot for *λ* tends to fall in between these two regimes.

Building an RPATH CV requires a careful process of selecting the portion of the system undergoing the conformational change, and choosing and optimising a number of reference structures. While structures of the end states are usually available, the intermediate structures have to be generated. One typically extrapolates them from biased simulations, such as steered MD or adiabatic bias MD^27,28^ (see Fig. 1A). These explorative simulations accelerate a reaction coordinate to quickly sample transitions connecting the end states, so that a trial sequence of conformations can be extracted from them. However, for large and complex biomolecular systems such as GPCRs, this process is particularly challenging and time-consuming, as it is difficult to both figure out a good reaction coordinate and to evaluate the quality of the extracted structures *a priori*. Eventually, preliminary biased simulations of the trial RPATH CV must be run to check whether the selected intermediates are sufficient to drive the desired conformational change. Numerous trial-and-error attempts are required before an effective CV is obtained. This recursive procedure is typically not transferable even between analogous systems, and the CV refinement process must be repeated for each new system under investigation.

**Figure 1:**
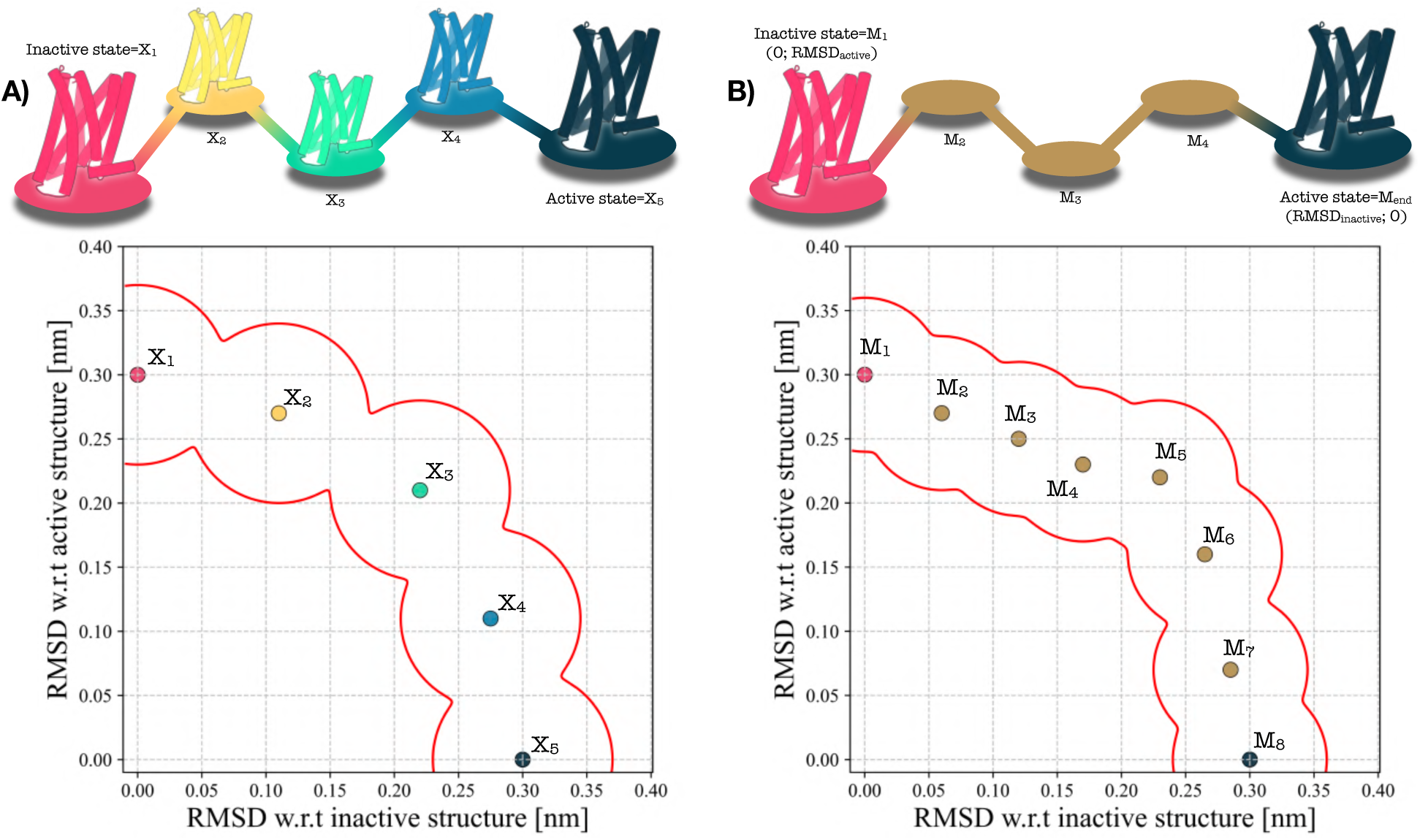
Schematic comparison between the original PATH CV ^26^ and the Euclidean variant used in this work. **a)** In the traditional formulation, constructing a meaningful PATH CV for GPCR activation requires multiple intermediate PDB structures *X_i_* along a putative transition pathway. Because these intermediates must be guessed and manually curated, the resulting path is laborious to set up and often uncertain in its direction, leading to repeated trial-and-error attempts. **b)** In the Euclidean variant, only the inactive and active structures are required, while all intermediate milestones are simply defined in terms of the two-dimensional space of RMSD values relative to these endpoints. This yields a path that is straightforward to define as it doesn’t depend on any intermediate structure.

A further limitation of RPATH CV is the requirement to focus the RMSD calculation on specific portions of the GPCR. Typically one has to align the RMSD over rigid portion of the GPCR and calculate the RMSD over the structural sections that change the most during the activation. Different alignment strategies on different sets of atoms yield vastly different RMSD measurements, critically impacting the quality of the resulting RPATH CV. This step as well requires a recursive optimisation process that further slows down the overall procedure. Although reformulating the path in terms of contact maps has been shown to mitigate this problem by avoiding explicit structural superposition^29–31^, a contact-map-based path CV remains difficult to build and cannot be transferred between systems.

A different and more straightforward approach was proposed to simulate the activation of the glucagon receptor^16^. The RMSDs over the two inactive and active end states were calculated and transformed into a CV by simply calculating their difference. The resulting CV captures by construction the progress along the GPCR activation and is equivalent to a linear path between two milestones. However, its lack of curvature represents a limit as it may force the system to pass through improbable high-energy states where both RMSDs are equally low. In the meantime, an alternative Euclidean formulation of the PATH CV, hereafter referred to as EPATH, was proposed and mostly used in the context of chemistry and catalysis^32–35^. In this variant, the path is not defined by the RMSD value with respect to a sequence of structures but as a progress in CV space. In other words, in EPATH the milestones *M_i_* are represented by tuples of CV values and do not need to be tied to real structures.

Here, we introduce a strategy that puts together these two approaches: we propose an EPATH CV on a two-dimensional space that uses the two RMSD values with respect to a GPCR’s inactive and active structures. A crucial component of our strategy is the availability of high resolution structures of the end states over which we measure the RMSD. Along the resulting EPATH CV, any visited structure of the system is encoded in a tuple *M* defined by a couple of RMSD values (*RMSD*_inactive_(*M*), *RMSD*_active_(*M*)). In this way, only the end states milestones, i.e., *M*_1_ and *M*_8_ in Fig. 1, have to be tied to real structures, while the milestones *M_i_* in between can be generated arbitrarily to chart a route in RMSD space. Here, we choose a slightly arched path above the diagonal that retraces the shape of the path that we observed in our previous work on GPCR activation^17^. This path connects the well defined end states by passing through an entropic basin where both RMSD values are high at the same time, that is, a basin populated by states that differ significantly from both the inactive and active references.

Within the EPATH CV approach, the Euclidean distance between the milestone value *M* of a state and any reference milestone *M_i_* is then simply defined as:

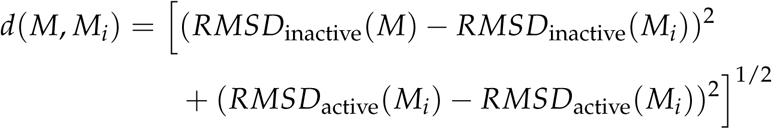

Substituting this distance into the expressions in Eq. 1 for *s*(*X*) and *z*(*X*) simply yields the EPATH CV value for structure *M* in a path made of N milestones:

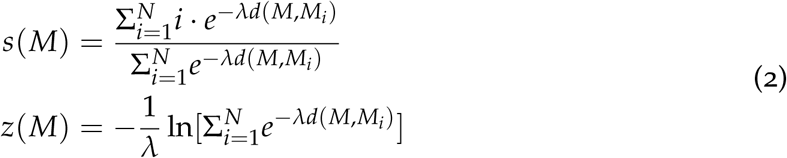

This approach retains the definitions of progress (*s*(*M*)) and deviation (*z*(*M*)) from the original RPATH CV approach. Its key advantage is that it avoids the need to define intermediate milestones tied to a curated set of conformations. Consequently, setting up the resulting EPATH CV is much simpler, as the time-consuming recursive refinement procedure of the previous approach is not required. Another advantage of this strategy is that it relies less on RMSD alignment and the selection of atoms used to calculate it. To highlight this, we choose to align and calculate the RMSD over the same set of C*α* atoms that make up the secondary structure of the system (more than 90% of the system’s atoms), and this generic choice had no need for refinement for building an efficient CV.

Finally, in terms of performance, the EPATH CV is faster to compute since the numerically expensive RMSD calculation needs to be carried out only twice per timestep. Conversely, in the RPATH CV such calculation is computed N times per timestep, once for each intermediate structural milestone. These advantages make applying the EPATH strategy to multiple systems more straightforward, reducing setup time and improving transferability.

### OneOPES MD simulations

The equilibrated structure of ADRB1 was taken from our previous work^17^ where we generated it starting from PDB ID: 7BVQ^36^. The structure of MOR was obtained from PDB ID: 9MQJ by removing the ligand resolved in the orthosteric binding site^37^. The details of the preparation and equilibration of MOR are detailed in SI and in Fig. S1.

We used OneOPES^25^, a replica-exchange extension of OPES Explore^38^, to sample the activation pathway of two GPCRs, ADRB1 and MOR. In each simulation, eight replicas were run, i.e., one convergence-focused (replica **0**) and seven exploratory ones (replicas **1–7**) where additional CVs were accelerated. All replicas shared OPES Explore as the main sampling bias, while replicas **1–7** were progressively heated up to 335 K by biasing the system’s potential energy with OPES Expanded^39^ to facilitate barrier crossing.

To directly compare the two approaches, we ran simulations using the GPCR activation PATH CV in both formulations, i.e., the original RPATH^26^ and the newer EPATH implementation. In analogy with Ref.^17^, additional CVs were introduced for four key microswitch motifs^40^, i.e., *PIF*, *DRY*, *NPxxY*, and *YY* to capture local structural determinants of activation. The distances within these motifs were biased as auxiliary CVs in the exploratory replicas. Both approaches use a similar set of parabolic restraints on the microswitches PIF and NPxxY to guide the system through the intermediate basins, which contains many metastable states and kinetic traps, to the fully active one. It is important to highlight that these parabolic restraints only apply to intermediates and not to end-point states, and as such they don’t affect their relative stability.

In addition, we biased the hydration of the cytoplasmic cavity, as our recent studies have shown that water coordination at specific sites can be critical for lowering the activation free-energy barrier^41–47^. In the case of ADRB1, this choice is further supported by NMR evidence indicating that the *YY*-motif is stabilized through a water-mediated interaction^48^.

For analyzing and comparing results, directly using the biased RPATH and EPATH CVs would be rather ineffective. We chose instead to reproject the free energy results on an alternative simplified EPATH CV with fewer milestones, six for ADRB1 and seven for MOR. Further analyses and figures on the biased PATH CVs are instead reported in the Supplementary Information (SI).

Simulations were performed using GROMACS 2023^49^ patched with PLUMED 2.9.1.^50,51^. Full details about simulation parameters and CVs are reported in the SI, respectively in Tables S1-S2 and Figures S2-S3, and the simulation input files are provided in GitHub at https://github.com/valeriorizzi/GPCR_Euclidean_PATH.git.

### Cluster analysis and Shannon’s entropy

We performed a cluster analysis on the MOR’s OneOPES simulations using GROMACS’s *gmx cluster* routine, using the *gromos* algorithm. An RMSD threshold value of 1.4 Å on the C*α*s was selected, considering the number of generated cluster families and the similarity of protein conformations within a cluster family. From the cluster population, the Shannon’s entropy was calculated, using the following formula:

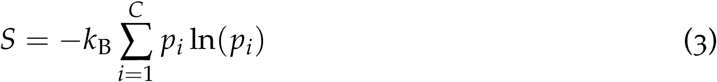

where *k*_B_ is the Boltzmann constant, *p_i_* is the fractional population of the *i*-th cluster *C* with respect to all the cluster families *C* and 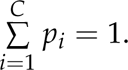 In the analysis, once we partition the phase space into different regions, we will use this metric as a natural measure of configurational diversity in each region. Regions with one dominant cluster family would display low entropy values, whereas regions with numerous equally probable cluster families would have a high entropy content.

## 3 Results

### Validation of the Euclidean PATH CV on ADRB1

We first assessed the performance of the EPATH CV on ADRB1, a prototypical class A GPCR that has been widely investigated both experimentally and computationally^36,48,52,53^. Using the same simulation setup and CV selection of our previous study^17^, we repeated OneOPES simulations of the apo-ADRB1 system, replacing the RPATH CV with the newly constructed one (see Fig. 2A and Fig. S4A). The resulting free-energy landscape along the activation pathway closely matches our earlier results, recovering both the relative stability of the inactive and active basins and the barrier separating them (see Fig. 2B). Notably, the simulations converge in significantly shorter timescales than with the RPATH CV, highlighting the efficiency gain of the EPATH approach without compromising accuracy (see Fig. 2C and Fig. S4B-C).

**Figure 2:**
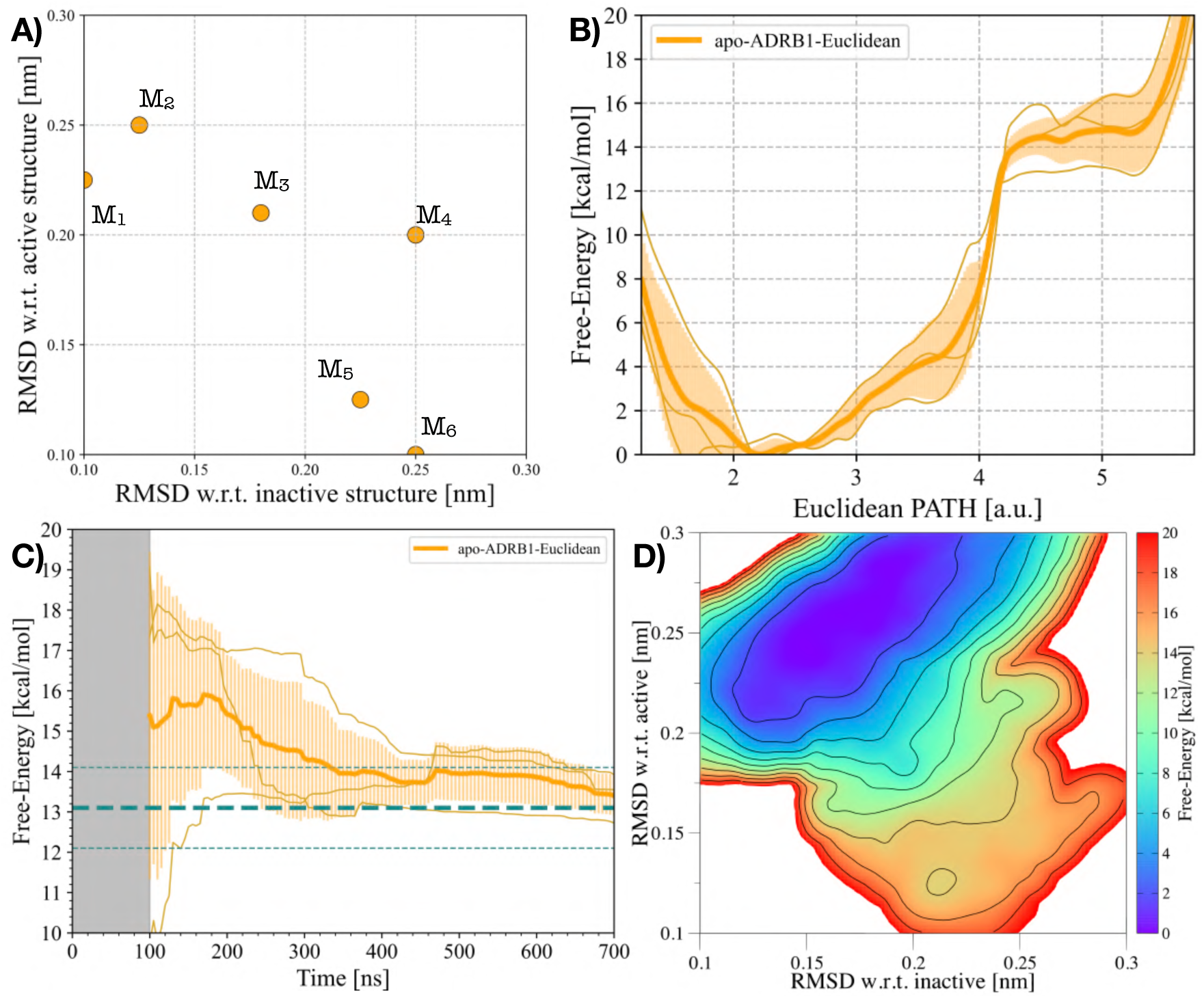
ADRB1’s activation using the EPATH CV. **a)** Milestones M*_i_*of the EPATH CV employed in *apo-ADRB1-Euclidean* and in Ref. ^17^ **b)** 1D FES as a function of the EPATH CV in (**a**) for the *apo-ADRB1-Euclidean* system. **c)** Free-energy difference between inactive and active states in the *apo-ADRB1-Euclidean* system over time. In (**b**) and (**c**), the average data over three replicas are represented through an orange solid line, while the transparent orange area illustrates the standard deviation. **d)** Average 2D FES concerning the RMSD of both the inactive and active structures for the *apo-ADRB1-Euclidean* system.

As a further form of comparison, we re-projected the accumulated bias potential onto the (*RMSD*_inactive_; *RMSD*_active_) plane, obtaining a two-dimensional free-energy surface (FES) describing the conformational change leading to ADRB1’s activation (Fig. 2D). This representation confirms that the 2D FES obtained with the EPATH is in excellent agreement with the one previously computed using the RPATH, with both inactive-and active-like basins recovered at the expected locations. Additional analyses of cavity hydration and the conformational rearrangements of key microswitches during ADRB1 activation are provided in the SI (see Fig. S5) and show a good quality agreement between the two approaches.

### Activation of the Mu Opioid Receptor

In this section, we report the simulation results of the activation of another class A GPCR: the *µ*-opioid receptor (MOR). MOR is one of the most critical analgesic targets and is at the centre of the ongoing opioid crisis^54,55^. We ran OneOPES with both the RPATH and the EPATH CV definition. In particular, the former has been extrapolated from a series of steered MD between representative inactive and active structures of MOR (for further details, refer to the SI and Fig. S6), while the latter has been defined through (*RMSD*_inactive_; *RMSD*_active_) tuples, following the same arched milestones distribution employed for ADRB1 (see Fig. 3A and S7). To deliver reliable statistics, each OneOPES simulation was carried out in triplicate. For clarity’s sake, the apo-MOR system sampled with the RPATH CV is referred to as *apo-MOR-OLD* hereafter, while the apo-MOR system sampled with the new EPATH CV as *apo-MOR-Euclidean*.

**Figure 3:**
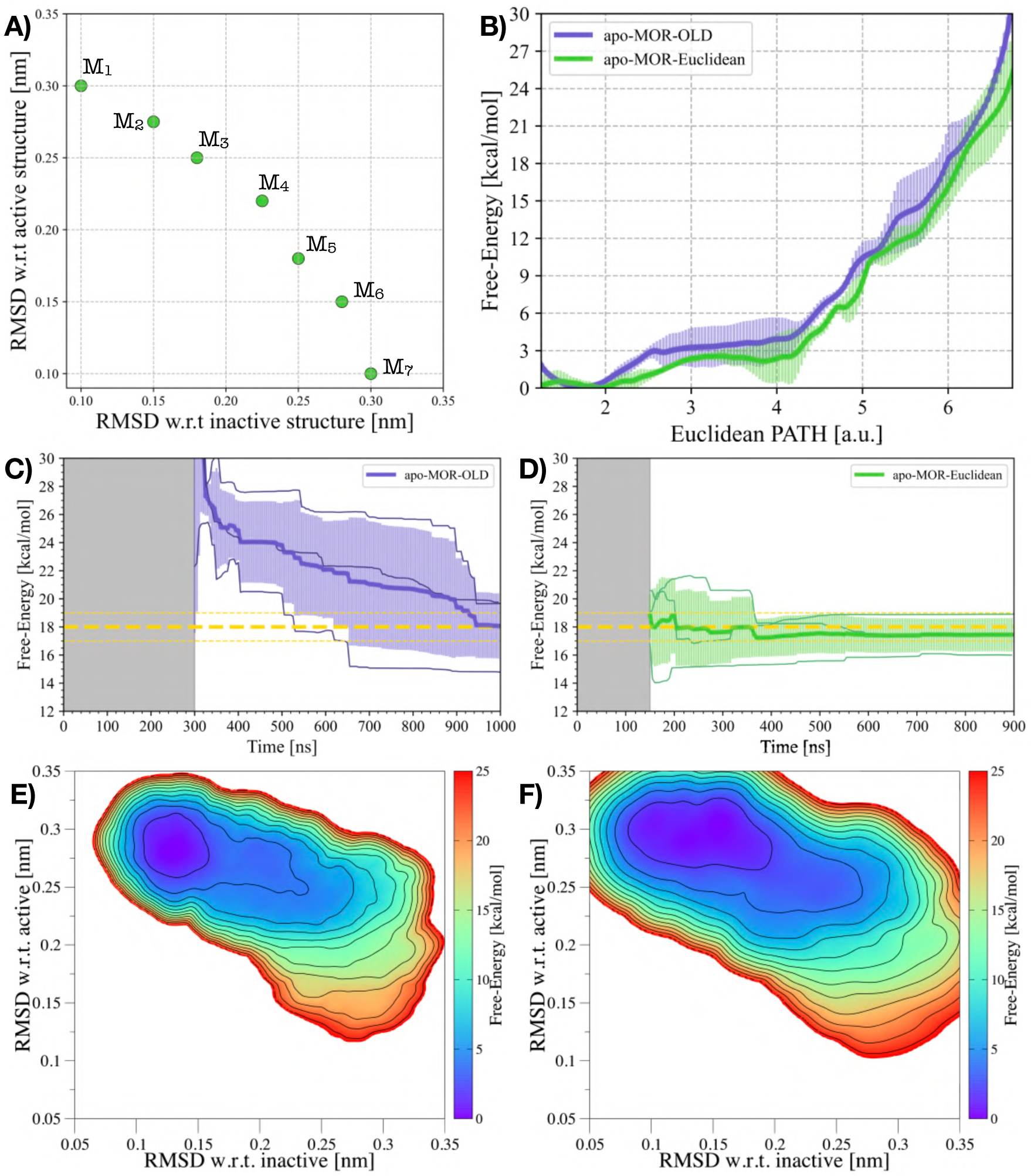
MOR’s activation using either the RPATH or the EPATH CVs. **a)** Milestones M*_i_* of the EPATH CV upon which the *apo-MOR-OLD* and *apo-MOR-Euclidean* free energies have been re-projected. **b)** Comparison of the 1D FES as a function of the EPATH CV in (**a**) for the apo-*apo-MOR-OLD* and *apo-MOR-Euclidean* systems. **c-d)** Free-energy difference between inactive and active states in the *apo-MOR-OLD* (**c**) and *apo-MOR-Euclidean* (**d**) systems over time. In (b) and (c), the average free energy difference of *apo-MOR-OLD* is represented by a purple solid line, while the transparent purple area illustrates the standard deviation. Similarly, in (b) and (d) the average free energy difference of *apo-MOR-OLD* is represented by a green solid line, while the transparent green area illustrates the standard deviation. **e-f)** Average 2D FES concerning the RMSD of both the inactive and active structures for the *apo-MOR-OLD* (**e**) and *apo-MOR-Euclidean* (**f**) systems.

The free-energy results obtained from the two approaches are in excellent agreement, with both capturing well the thermodynamic equilibrium between inactive- and active-like states. The agreement is evident from the one-dimensional free energy obtained by re-weighting the accumulated bias potential of *apo-MOR-OLD* onto *apo-MOR-Euclidean*’s EPATH CV (see Fig. 3B). However, the RPATH CV takes a rather long time to converge with the free energy difference reaching an agreement between the independent replicas only at the end of the simulation (see Fig. 3C). This indicates that, even though the CV itself went through an iterative optimization process, it is still far from ideal, and further refinements are necessary.

On the contrary, the EPATH CV once again converges faster than the RPATH, requiring a way shorter simulation time to achieve comparable estimates of the free-energy difference among the three independent replicas (see Fig. 3D). Remarkably, no recursive CV optimization was needed to refine this PATH CV. In details, for *apo-MOR-OLD*, the ΔG of activation is estimated to be 18.0 *±* 1.0 kcal/mol after *∼* 800 ns, whereas *apo-MOR-Euclidean*’s OneOPES simulations converge to the ΔG value of *∼*18.0 kcal/mol after only 200 ns. Similar considerations apply to the 2D FES measured on the (*RMSD*_inactive_; *RMSD*_active_) space: both sets of simulations are able to explore an analogous conformational space (see Fig. 3E and 3F), with the *apo-MOR-Euclidean* trajectory being able to sample more thoroughly the fully active basin (*RMSD*_active_ < 0.1 nm, see Fig. S7 for more details).

To further assess and compare the sampling quality of the two approaches, we additionally analyzed the rearrangement of key MOR microswitches along the activation coordinate. Specifically, we computed 2D FESs as a function of the activation pathway and of structural descriptors associated with the *NPxxY*, *DRY*, and *YY* motifs. In all cases, the *apo-MOR-OLD* and *apo-MOR-Euclidean* simulations delivered similar landscapes, capturing both the qualitative ordering of microswitch transitions and the quantitative positioning of the free-energy minima (see Fig. S8). We also calculated a 2D FES to characterize the hydration dynamics of the intracellular cavity in proximity of the *YY* motif, a well-known hallmark of GPCR activation. This analysis reveals again a strong agreement between the two PATH CV definitions, with both approaches identifying analogous hydration patterns and their coupling to the receptor’s progression along the activation pathway (Fig. S9).

The quality of convergence and agreement between approaches across two distinct GPCR systems further supports that the EPATH CV can reliably replace the traditional trial-and-error workflow from the RPATH.

### Advantages of the Euclidean PATH

To better understand how EPATH effectively drives GPCR activation, we analyzed the distribution of sampled configurations across the two collective variables, *s*(*M*) and *z*(*M*). In EPATH, the intermediate values of the *s*(*M*) coordinate correspond to conformations that are simultaneously distant from both the inactive and active references in RMSD space. It is worth stressing that RMSD is a spherical yet anisotropic descriptor: a high RMSD value corresponds to a multitude of structurally distinct states equally distant from the reference state^56^. For this reason, using RMSD alone as a CV in enhanced sampling is not particularly effective and may lead to poorly converged results.

In contrast, the EPATH CV couples two RMSD measures, thereby embedding directionality throughout its range in a push-pull fashion. This bivariate description enables detours away from the end states, while avoiding the vast and uninformative region of high-RMSD configurations where simulations typically suffer from hysteresis. An important role in this context is played by the presence of loose restraints on the path deviation *z* and of coupled restraints between the progress CV *s* and microswitches *PIF* and *NPxxY*. This helps the convergence by aligning the conformation of the microswitches with the activation of the macroswitches. Another important element is the presence of the membrane that confines the GPCR and prevents it from exploring high-in-energy partially unfolded states.

Importantly, EPATH does not require a tedious definition of all the intermediate milestones. Instead, in RPATH the definition of these milestones is crucial. The milestones are not known *a priori* and are typically extracted from steered MD trajectories. A sub-optimal choice of the intermediate milestones decreases the efficiency of the method by forcing the sampling of conformations that are far from the RPATH. EPATH has fine resolution near the endpoints (where the corresponding structures and energy basins are well defined), while remaining less well defined in the middle, where the receptor naturally explores a broader ensemble. When combined with OneOPES, and auxiliary biases on the microswitches and water, this transition state region is crossed rapidly and reversibly, ensuring quicker convergence than with RPATH.

This behaviour is clearly illustrated in Fig. 4. In both *apo-MOR-OLD* (see Fig. 4A) and *apo-MOR–Euclidean* (see Fig. 4B), the intermediate portions of the EPATH are characterized by a plethora of slightly out-of-path frames (mostly between *s*(*M*) *∼* 4 and *s*(*M*) *∼* 5). Performing a cluster analysis on the trajectory frames confirms that the intermediate portion of the EPATH encompasses distinct structural families (see Fig. S10 and S11 for further information). Notably, the simulation frames belonging to the first and last milestones (i.e., *M*_1_ and *M*_7_) can be grouped in very few cluster families, with the most probable ones containing more than 95% of the cluster populations. Conversely, the more we approach the central values of the EPATH, the more we can appreciate both an increase in cluster families and a decrease in their relative abundance.

**Figure 4:**
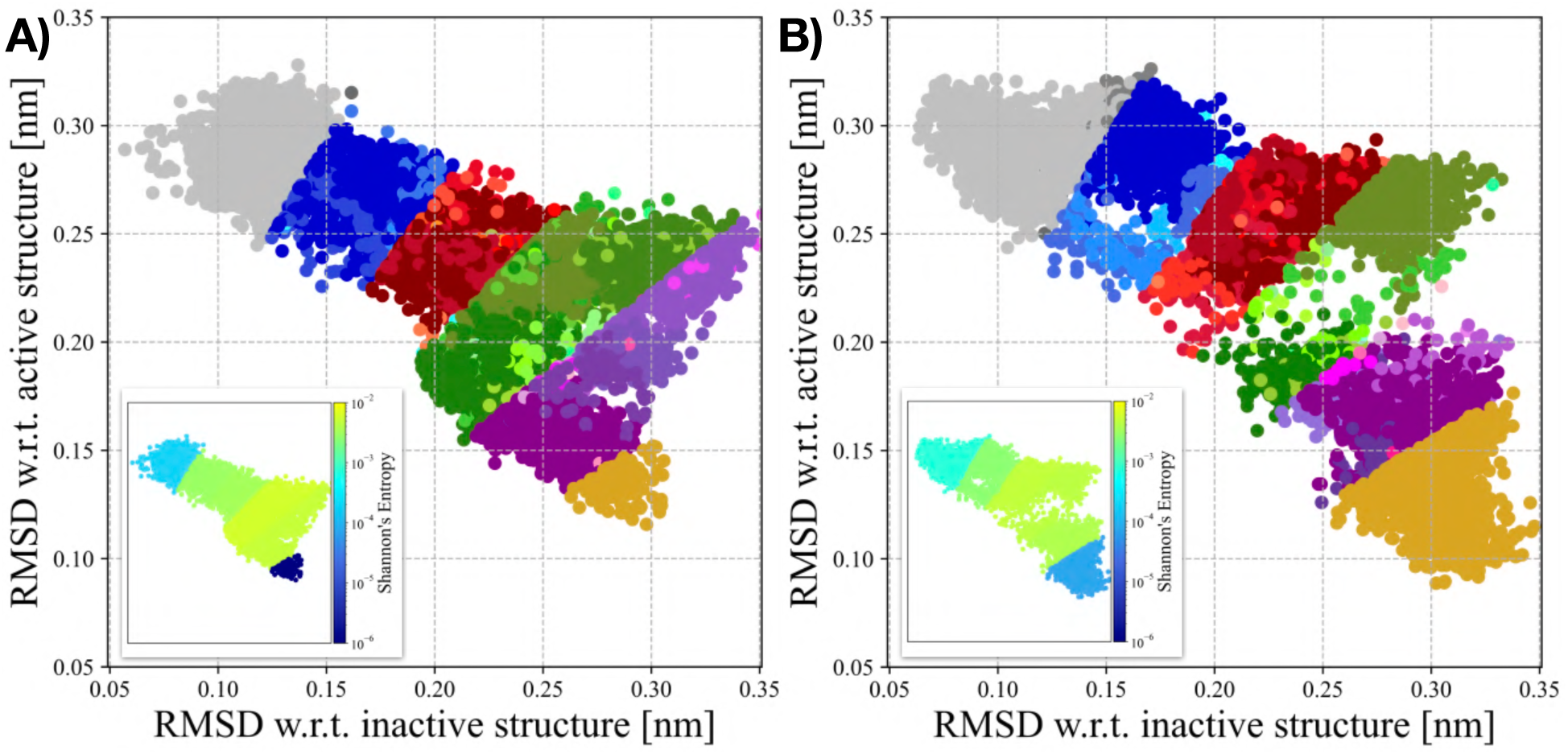
Analysis of the conformational heterogeneity observed in *apo-MOR-OLD* and *apo-MOR-Euclidean* OneOPES simulations. **a-b)** Sampling distributions of the *apo-MOR-OLD* (**a**) and *apo-MOR-Euclidean* (**b**) OneOPES simulations projected onto the (*RMSD*_inactive_; *RMSD*_active_) space. Frames are grouped according to their Euclidean PATH interval (M1–M2, M2–M3, M3–M4, M4–M5, M5–M6, M6–M7), and points within each interval are coloured according to the cluster they belong to (grey, blue, red, green, purple, and yellow palettes, respectively). In both panels, a small inset displays the same (*RMSD*_inactive_; *RMSD*_active_) sampling coloured by the local Shannon entropy, providing a complementary view of the conformational heterogeneity within each region of the activation landscape.

To quantitatively characterize how the size of the conformational pool varies along the progress coordinate, we turned to information theory and computed the Shannon entropy associated with the cluster populations at each milestone (see “Methods" for additional details). Consistent with our qualitative observations, both *apo-MOR-OLD* and *apo-MOR-Euclidean* exhibit the same global behaviour: entropy reaches its minimum at the terminal milestones, where the conformational space is tightly focused around well-defined inactive and active structures, and rises sharply toward the center of the path (see Fig. 4A-B). Remarkably, in both formulations, the entropy in the *M*_4_ *− M*_5_ interval increases by nearly three orders of magnitude relative to the endpoints, quantitatively confirming that the intermediate milestones host a vastly broader and more diverse ensemble of receptor conformations. This information-theoretic analysis reinforces the picture emerging from clustering and structural projections: the EPATH naturally captures the physical widening of the conformational landscape in the transition region, while maintaining strong definition at the two endpoints.

## 4 Conclusions

GPCRs are central regulators of cellular signalling and remain the targets of a large fraction of therapeutic compounds. A key determinant of their pharmacological action is the balance between inactive and active conformational states, which underlies the agonist–antagonist axis. Obtaining reliable free-energy profiles for these transitions is therefore a critical step toward understanding receptor function and quantifying the impact of ligand binding. However, the inherent complexity of GPCR conformational landscapes poses major challenges for molecular simulations, requiring strategies that are accurate, practical and transportable^8,57^.

In this work, we introduced a streamlined protocol for building PATH CVs within the OneOPES enhanced sampling framework. By eliminating the tedious trial-and-error procedure of standard path definition, the new approach reduces the user effort required to set up simulations while maintaining the accuracy of the computed free-energy landscapes. Its application to two prototypical receptors^58^, ADRB1 and MOR, demonstrates that this approach reproduces the thermodynamic results of the traditional method with a significantly faster convergence. An in-depth analysis of MOR further confirmed that the mechanistic details of activation, including side-chain microswitches and hydration changes, are faithfully captured.

An important question is whether providing additional information about the intermediate region of the activation landscape could further improve the approach. In our view, a balance must be struck between ease of use and computational efficiency. For in-depth mechanistic studies of a single receptor, more elaborate CV refinement strategies may indeed offer advantages. However, when the goal is to analyse multiple GPCRs and to reliably estimate activation free energies without extensive system-specific optimisation, the Euclidean PATH approach presented in this manuscript provides a robust, easy-to-build solution that delivers reliable sampling between end states.

Nonetheless, it is also important to acknowledge the limitations of RMSD-based metrics. As previously stated, at large RMSD values the number of distinct conformations grows rapidly, making such regions difficult to sample and ineffective to bias. In our opinion, the success of the present approach is largely due to the constrained conformational space imposed by the mild restraining potentials imposed on the intermediate states assumed by the microswitches and the membrane environment, which limits the overall extent of RMSD exploration. Systems that experience broader structural re-arrangement embedded in a less constrained environment, e.g., a globular protein in water solution, may demand additional care, as the effective size of RMSD space increases and sampling may become a major challenge. Another point to bear in mind is that the estimated barriers under these conditions will not be as reliable as the difference between the end-point (inactive and active) states.

Overall, this methodological refinement represents an important step toward the routine use of enhanced sampling approaches to study GPCRs. By improving both efficiency and accessibility, it paves the way for future investigations that extend beyond the apo receptors considered here to quantify the effect of ligands on activation free energies. Ultimately, our proposed approach will facilitate a more systematic assessment of how ligands affect GPCR conformational equilibria, providing valuable guidance for drug discovery and pharmacological design.

## Data Availability Statement

The OneOPES enhanced sampling simulation input files are available on GitHub at the link https://github.com/valeriorizzi/GPCR_Euclidean_PATH.git. The script requires PLUMED^50,51^, version 2.8 or later. The enhanced sampling simulations are run with GROMACS 2023^49^.

## Supporting Information Available

Additional figures, tables and computational details on the CV building strategy and on OneOPES enhanced sampling simulations.

## Acknowledgement

This work was financially supported by the Swiss National Science Foundation and Bridge funding schemes (project numbers: 200021_204795, CRSII5_216587, and 40B2-0_203628). We acknowledge the Swiss National Supercomputing Centre for supercomputer time allocations on Piz Daint and Alps (project ID: s1274, and lp84). We wish to thank Gareth Tribello and Alberto Borsatto for useful discussions.

## References

(1) Rask-Andersen, M.; Almén, M. S.; Schiöth, H. B. Trends in the exploitation of novel drug targets. Nature reviews Drug discovery 2011, 10, 579–590.

(2) Katritch, V.; Cherezov, V.; Stevens, R. C. Structure-function of the G protein–coupled receptor superfamily. Annual review of pharmacology and toxicology 2013, 53, 531–556.

(3) Sriram, K.; Insel, P. A. G protein-coupled receptors as targets for approved drugs: how many targets and how many drugs? Molecular pharmacology 2018, 93, 251–258.

(4) Weis, W. I.; Kobilka, B. K. The molecular basis of G protein–coupled receptor activation. Annual review of biochemistry 2018, 87, 897–919.

(5) Venkatakrishnan, A.; Deupi, X.; Lebon, G.; Tate, C. G.; Schertler, G. F.; Babu, M. M. Molecular signatures of G-protein-coupled receptors. Nature 2013, 494, 185–194.

(6) Dror, R. O.; Green, H. F.; Valant, C.; Borhani, D. W.; Valcourt, J. R.; Pan, A. C.; Arlow, D. H.; Canals, M.; Lane, J. R.; Rahmani, R. et al. Structural basis for modulation of a G-protein-coupled receptor by allosteric drugs. Nature 2013, 503, 295–299.

(7) Huang, W.; Manglik, A.; Venkatakrishnan, A. J.; Laeremans, T.; Feinberg, E. N.; Sanborn, A. L.; Kato, H. E.; Livingston, K. E.; Thorsen, T. S.; Kling, R. C. et al. Structural insights into µ-opioid receptor activation. Nature 2015, 524, 315–321.

(8) Latorraca, N. R.; Venkatakrishnan, A. J.; Dror, R. O. GPCR Dynamics: Structures in Motion. Chem. Rev. 2017, 117, 139–155.

(9) Lopez-Balastegui, M.; Stepniewski, T. M.; Kogut-Günthel, M. M.; Di Pizio, A.; Rosenkilde, M. M.; Mao, J.; Selent, J. Relevance of G protein-coupled receptor (GPCR) dynamics for receptor activation, signalling bias and allosteric modulation. British Journal of Pharmacology 2024, n/a, 1–14, _eprint: https://onlinelibrary.wiley.com/doi/pdf/10.1111/bph.16495.

(10) De Felice, A.; Aureli, S.; Limongelli, V. Drug repurposing on G protein-coupled receptors using a computational profiling approach. Frontiers in molecular biosciences 2021, 8, 673053.

(11) Aranda-García, D.; Stepniewski, T. M.; Torrens-Fontanals, M.; García-Recio, A.; Lopez-Balastegui, M.; Medel-Lacruz, B.; Morales-Pastor, A.; Peralta-García, A.; Dieguez-Eceolaza, M.; Sotillo-Nuñez, D. et al. Large scale investigation of GPCR molecular dynamics data uncovers allosteric sites and lateral gateways. Nat Commun 2025, 16, 2020.

(12) Calderón, J. C.; Ibrahim, P.; Gobbo, D.; Gervasio, F. L.; Clark, T. Activa-tion/Deactivation Free-Energy Profiles for the *β*2-Adrenergic Receptor: Ligand Modes of Action. Journal of Chemical Information and Modeling 2023, 63, 6332–6343.

(13) Conflitti, P.; Lyman, E.; Sansom, M. S. P.; Hildebrand, P. W.; Gutiérrez-de Terán, H.; Carloni, P.; Ansell, T. B.; Yuan, S.; Barth, P.; Robinson, A. S. et al. Functional dynamics of G protein-coupled receptors reveal new routes for drug discovery. Nat Rev Drug Discov 2025, 1–25, Publisher: Nature Publishing Group.

(14) D’Amore, V. M.; Conflitti, P.; Marinelli, L.; Limongelli, V. Minute-timescale free-energy calculations reveal a pseudo-active state in the adenosine A2A receptor activation mechanism. Chem 2024, 10, 3678–3698, Publisher: Elsevier.

(15) Provasi, D.; Artacho, M. C.; Negri, A.; Mobarec, J. C.; Filizola, M. Ligand-induced modulation of the free-energy landscape of G protein-coupled receptors explored by adaptive biasing techniques. PLoS Comput Biol 2011, 7, e1002193.

(16) Mattedi, G.; Acosta-Gutiérrez, S.; Clark, T.; Gervasio, F. L. A combined activation mechanism for the glucagon receptor. Proceedings of the National Academy of Sciences 2020, 117, 15414–15422.

(17) Aureli, S.; Rizzi, V.; Piasentin, N.; Gervasio, F. L. Enhanced sampling and tailored collective variables yield reproducible free energy landscapes of beta-1 adrenergic receptor activation. Journal of Chemical Theory and Computation 2025, 21, 7687–7700.

(18) Rodríguez-Espigares, I.; Torrens-Fontanals, M.; Tiemann, J. K. S.; Aranda-García, D.; Ramírez-Anguita, J. M.; Stepniewski, T. M.; Worp, N.; Varela-Rial, A.; Morales-Pastor, A.; Medel-Lacruz, B. et al. GPCRmd uncovers the dynamics of the 3D-GPCRome. Nat Methods 2020, 17, 777–787.

(19) Calderón, J. C.; Ibrahim, P.; Gobbo, D.; Gervasio, F. L.; Clark, T. General Metadynamics Protocol To Simulate Activation/Deactivation of Class A GPCRs: Proof of Principle for the Serotonin Receptor. J. Chem. Inf. Model. 2023, 63, 3105–3117, Publisher: American Chemical Society.

(20) Saleh, N.; Saladino, G.; Gervasio, F. L.; Clark, T. Investigating allosteric effects on the functional dynamics of b2-adrenergic ternary complexes with enhanced-sampling simulations. Chemical science 2017, 8, 4019–4026.

(21) Hénin, J.; Lelièvre, T.; Shirts, M. R.; Valsson, O.; Delemotte, L. Enhanced Sampling Methods for Molecular Dynamics Simulations [Article v1.0]. Living Journal of Computational Molecular Science 2022, 4, 1583–1583, Number: 1.

(22) Laio, A.; Gervasio, F. L. Metadynamics: a method to simulate rare events and reconstruct the free energy in biophysics, chemistry and material science. Reports on Progress in Physics 2008, 71, 126601.

(23) Fröhlking, T.; Aureli, S.; Gervasio, F. L. Learning committor-consistent collective variables: Transition pathways. Nature Computational Science 2025, 5, 520–521.

(24) Yuan, X.; Raniolo, S.; Limongelli, V.; Xu, Y. The molecular mechanism underlying ligand binding to the membrane-embedded site of a G-protein-coupled receptor. Journal of chemical theory and computation 2018, 14, 2761–2770.

(25) Rizzi, V.; Aureli, S.; Ansari, N.; Gervasio, F. L. OneOPES, a combined enhanced sampling method to rule them all. Journal of Chemical Theory and Computation 2023, 19, 5731–5742.

(26) Branduardi, D.; Gervasio, F. L.; Parrinello, M. From A to B in free energy space. The Journal of chemical physics 2007, 126, 054103.

(27) Marchi, M.; Ballone, P. Adiabatic bias molecular dynamics: a method to navigate the conformational space of complex molecular systems. The Journal of chemical physics 1999, 110, 3697–3702.

(28) Park, S.; Schulten, K. Calculating potentials of mean force from steered molecular dynamics simulations. The Journal of chemical physics 2004, 120, 5946–5961.

(29) Sutto, L.; Gervasio, F. L. Effects of oncogenic mutations on the conformational free-energy landscape of EGFR kinase. Proceedings of the National Academy of Sciences 2013, 110, 10616–10621.

(30) Kuzmanic, A.; Sutto, L.; Saladino, G.; Nebreda, A. R.; Gervasio, F. L.; Orozco, M. Changes in the free-energy landscape of p38*α* MAP kinase through its canonical activation and binding events as studied by enhanced molecular dynamics simulations. eLife 2017, 6.

(31) Meral, D.; Provasi, D.; Filizola, M. An efficient strategy to estimate thermodynamics and kinetics of G protein-coupled receptor activation using metadynamics and maximum caliber. The Journal of chemical physics 2018, 149, 224101.

(32) Pietrucci, F.; Saitta, A. M. Formamide reaction network in gas phase and solution via a unified theoretical approach: Toward a reconciliation of different prebiotic scenarios. Proceedings of the National Academy of Sciences 2015, 112, 15030–15035.

(33) Polino, D.; Parrinello, M. Kinetics of Aqueous Media Reactions via Ab Initio Enhanced Molecular Dynamics: The Case of Urea Decomposition. The Journal of Physical Chemistry B 2019, 123, 6851–6856.

(34) Das, S.; Raucci, U.; Neves, R. P. P.; Ramos, M. J.; Parrinello, M. How and When Does an Enzyme React? Unraveling *α*-Amylase Catalytic Activity with Enhanced Sampling Techniques. ACS Catalysis 2023, 13, 8092–8098.

(35) Das, S.; Raucci, U.; Neves, R. P. P.; Ramos, M. J.; Parrinello, M. Correlating enzymatic reactivity for different substrates using transferable data-driven collective variables. Proceedings of the National Academy of Sciences 2024, 121, 2017.

(36) Xu, X.; Kaindl, J.; Clark, M. J.; Hübner, H.; Hirata, K.; Sunahara, R. K.; Gmeiner, P.; Kobilka, B. K.; Liu, X. Binding pathway determines norepinephrine selectivity for the human *β*1AR over *β*2AR. Cell Research 2021, 31, 569–579.

(37) Vigneron, S. F.; Ohno, S.; Braz, J.; Kim, J. Y.; Kweon, O. S.; Webb, C.; Billesbølle, C. B.; Srinivasan, K.; Bhardwaj, K.; Irwin, J. J. et al. Docking 14 Million Virtual Isoquinuclidines against the *µ* and *κ* Opioid Receptors Reveals Dual Antagonists–Inverse Agonists with Reduced Withdrawal Effects. ACS Central Science 2025, 11, 770–790.

(38) Invernizzi, M.; Parrinello, M. Exploration vs convergence speed in adaptive-bias enhanced sampling. Journal of Chemical Theory and Computation 2022, 18, 3988–3996.

(39) Invernizzi, M.; Piaggi, P. M.; Parrinello, M. Unified approach to enhanced sampling. Physical Review X 2020, 10, 041034.

(40) Hauser, A. S.; Kooistra, A. J.; Munk, C.; Heydenreich, F. M.; Veprintsev, D. B.; Bouvier, M.; Babu, M. M.; Gloriam, D. E. GPCR activation mechanisms across classes and macro/microscales. Nature structural & molecular biology 2021, 28, 879–888.

(41) Rizzi, V.; Bonati, L.; Ansari, N.; Parrinello, M. The role of water in host-guest interaction. Nature Communications 2021, 12, 93.

(42) Ansari, N.; Rizzi, V.; Parrinello, M. Water regulates the residence time of Benzamidine in Trypsin. Nature Communications 2022, 13, 5438.

(43) Karrenbrock, M.; Borsatto, A.; Rizzi, V.; Lukauskis, D.; Aureli, S.; Luigi Gervasio, F. Absolute Binding Free Energies with OneOPES. The Journal of Physical Chemistry Letters 2024, 15, 9871–9880.

(44) Ding, X.; Aureli, S.; Vaithia, A.; Lavriha, P.; Schuster, D.; Khanppnavar, B.; Li, X.; Blum, T. B.; Picotti, P.; Gervasio, F. L. et al. Structural basis of connexin-36 gap junction channel inhibition. Cell Discovery 2024, 10, 68.

(45) Febrer Martinez, P.; Rizzi, V.; Aureli, S.; Gervasio, F. L. Host–Guest Binding Free Energies à la Carte: An Automated OneOPES Protocol. Journal of Chemical Theory and Computation 2024, 10275–10287.

(46) Aureli, S.; Bellina, F.; Rizzi, V.; Gervasio, F. L. Investigating ligand-mediated conformational dynamics of pre-miR21: A machine-learning-aided enhanced sampling study. Journal of Chemical Information and Modeling 2024, 64, 8595–8603.

(47) Schulze, M.; Khakhula, T.; Piasentin, N.; Aureli, S.; Rizzi, V.; Gervasio, F. L. All you need is water: Converging ligand binding simulations with hydration collective variables. The Journal of Chemical Physics 2025, 163.

(48) Grahl, A.; Abiko, L. A.; Isogai, S.; Sharpe, T.; Grzesiek, S. A high-resolution description of *β*1-adrenergic receptor functional dynamics and allosteric coupling from backbone NMR. Nature communications 2020, 11, 2216.

(49) Abraham, M. J.; Murtola, T.; Schulz, R.; Páll, S.; Smith, J. C.; Hess, B.; Lindahl, E. GROMACS: High performance molecular simulations through multi-level parallelism from laptops to supercomputers. SoftwareX 2015, 1, 19–25.

(50) Tribello, G. A.; Bonomi, M.; Branduardi, D.; Camilloni, C.; Bussi, G. PLUMED 2: New feathers for an old bird. Computer physics communications 2014, 185, 604–613.

(51) Tribello, G. A.; Bonomi, M.; Bussi, G.; Camilloni, C.; Armstrong, B. I.; Arsiccio, A.; Aureli, S.; Ballabio, F.; Bernetti, M.; Bonati, L. et al. PLUMED Tutorials: A collaborative, community-driven learning ecosystem. The Journal of Chemical Physics 2025, 162.

(52) Abiko, L. A.; Grahl, A.; Grzesiek, S. High pressure shifts the *β*1-adrenergic receptor to the active conformation in the absence of G protein. Journal of the American Chemical Society 2019, 141, 16663–16670.

(53) Su, M.; Paknejad, N.; Zhu, L.; Wang, J.; Do, H. N.; Miao, Y.; Liu, W.; Hite, R. K.; Huang, X.-Y. Structures of *β*1-adrenergic receptor in complex with Gs and ligands of different efficacies. Nature communications 2022, 13, 4095.

(54) Volkow, N. D.; Blanco, C. The changing opioid crisis: development, challenges and opportunities. Molecular psychiatry 2021, 26, 218–233.

(55) Che, T.; Roth, B. L. Molecular basis of opioid receptor signaling. Cell 2023, 186, 5203–5219.

(56) Kufareva, I.; Abagyan, R. In Homology Modeling: Methods and Protocols; Orry, A. J. W., Abagyan, R., Eds.; Humana Press: Totowa, NJ, 2012; pp 231–257.

(57) Anantakrishnan, S.; Naganathan, A. N. Thermodynamic architecture and conformational plasticity of GPCRs. Nature Communications 2023, 14, 128.

(58) Marino, K. A.; Prada-Gracia, D.; Provasi, D.; Filizola, M. Impact of lipid composition and receptor conformation on the spatiotemporal organization of *µ*-opioid receptors in a multi-component plasma membrane model. PLoS computational biology 2016, 12, e1005240.

