## Supplementary Material for "A Transferable and Robust Computational Framework for Class A GPCR Activation Free Energies"

The data hereby reported are divided into the following sections:

- Supplementary Data 1: Systems preparation;
- Supplementary Data 2: Details on the employed CVs and how they have been developed;
- Supplementary Data 3: Additional data on the apo-ADRB1-Euclidean OneOPES simulation;
- Supplementary Data 4: Details on apo-ADRB1's microswitches and hydration during the activation;
- Supplementary Data 5: Frameworks of the apo-MOR-OLD and apo-MOR-Euclidean OneOPES simulations;
- Supplementary Data 6: Microswitches' behaviour of MOR during the GPCR activation;
- Supplementary Data 7: Cluster analysis of *apo-MOR-OLD* and *apo-MOR-Euclidean* OneOPES simulations.

### Supplementary Data 1

#### Preparation of MOR system

The structure of MOR was obtained from PDB ID: 9MQJ by removing the co-crystallized ligand [1]. The residues $H^{45.56}$ and $P^{5.30}$ are missing in the 9MQJ structure and were modelled with ChimeraX [2] to complete the receptor. A disulfide bridge was introduced between $C^{3.25}$ and $C^{45.50}$. To properly simulate the wild-type human MOR, ChimeraX's rotamer library [3] was exploited to reconstruct the side-chains of 54 9MQJ's amino acids, which were incomplete in the original PDB (see Fig. S1A-B).

A)

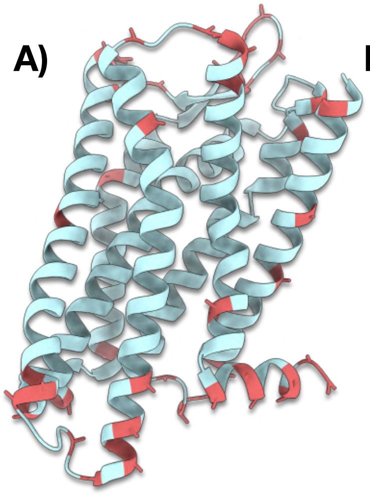

B)

| TM1 | ICL1 | TM2 | ECL1 | TM3 | ICL2 | TM4 | ECL2 | TM5 | ICL3 | TM6 | ECL3 | TM7 | H8 |
|---|---|---|---|---|---|---|---|---|---|---|---|---|---|
| 78 | 110 | 139 | | | 186 | 195 | 222 | 239 | 279 | 281 | 320 | 324 | 354 |
| 80 | 111 | 142 | | | 187 | 211 | 223 | 242 | | 283 | 323 | 325 | 357 |
| 90 | 113 | | | | 189 | | 224 | 256 | | 284 | | 328 | 358 |
| 97 | | | | | 190 | | 225 | 271 | | 286 | | 348 | 361 |
| 103 | | | | | | | 227 | 273 | | 290 | | | 362 |
| 107 | | | | | | | 229 | 274 | | 312 | | | 364 |
| 108 | | | | | | | 233 | | | | | | 365 |
| 109 | | | | | | | 235 | | | | | | |
| | | | | | | | 238 | | | | | | |

Figure S1: **Identification of unresolved side-chain residues in the MOR structure. a)** Mapping of all MOR residues with unresolved side-chain atoms in the cryo-EM structure PDB ID 9MQJ (coloured in cyan). Residues in red indicate positions where side-chain coordinates are missing in the deposited model, potentially affecting local interactions or structural interpretations; **b)** Summary table listing these unresolved residues, organised by the corresponding secondary-structure elements of MOR (e.g., transmembrane helices, intracellular loops, extracellular loops).

The mended MOR receptor was embedded into a membrane model (POPC/CHL 80:20) using CHARMM-GUI [4], and solvated with the TIP4PD water model and a salinity of 0.15 M NaCl. The N-terminus and C-terminus of the MOR structure were capped with acetyl and methyl-amino protecting groups, respectively. The DES-Amber force field was employed [5] in the MD engine GROMACS 2023 [6]. To equilibrate the system, a thermalisation cycle was applied with progressively reduced restraints on heavy atoms to relax any structural artifacts while maintaining the global organisation of the protein and membrane. The cycle consists of successive 1 ns NVT and 1 ns NPT simulations at increasing temperatures from 100 to 300 K in 50 K increments. Hydrogen bonds were constrained with LINCS [7, 8], and an integration step of 2 fs was used. Coulomb and van der Waals interactions were truncated at 1.0 nm, and long-range electrostatics were treated with the particle–mesh Ewald (PME) method [9]. The temperature was set at 300 K and controlled with the V-rescale thermostat [10], whereas the pressure was fixed at a reference value of 1 bar with the semi-isotropic C-rescale barostat [11].

#### Preparation of ADRB1 system

The structure of ADRB1 was taken from our previous work [12], generated starting from PDB ID: 7BVQ [13]. As discussed in Ref. [12], ADRB1's inactive structure encompasses only a portion of the whole human ADRB1 sequence, notably residues $S^{1.28}$ to $S^{5.74}$ and $V^{6.26}$ to $C^{8.59}$. Moreover, the ends of the sequences $S^{5.74}$ and $V^{6.26}$ were artificially linked, resulting in a shorter intracellular loop 3 compared to the native sequence of human ADRB1. As described in previous studies [13], the cysteine couples $C^{3.25}$-$C^{45.50}$

and $C^{45.43}$-$C^{45.49}$ were bound with a disulfide bridge, whereas $E^{3.41}$ was protonated as it predominantly exists in its neutral form. Protecting groups at ADRB1's termini, membrane composition, and salinity are the same as those used in our previous publication[12]. The ADRB1 simulations were performed using the same force field and simulation protocol as described for MOR, including thermostat, barostat, integration step, and cutoff settings.

### Supplementary Data 2

To investigate the activation pathway of the MOR GPCR in both the *apo-MOR-OLD* and *apo-MOR-Euclidean* systems, we employed a set of dedicated CVs reported in Tab. S1 and S2, respectively. These variables are specifically designed to effectively capture critical structural and hydration-related changes that occur during receptor activation, highlighting the complexity and significance of this vital biological process. As in Ref [12], the CVs on the residue couples (i.e., **D1-D7**) were built following a coarse-grained-inspired approach, treating the entire side-chain of specific amino acids as a putative dummy atom. For a detailed atomistic description of the **D1-D7** CVs, please refer to Fig. S2. Regarding the EPATH CV (i.e., Neop1.s) displayed in Fig. 3A, we reported a blueprint of the PLUMED file necessary to run it in Fig. S3.

| Replicas | 0 | 1 | 2 | 3 | 4 | 5 | 6 | 7 |
|---|---|---|---|---|---|---|---|---|
| OPES Explore | p1.s | p1.s | p1.s | p1.s | p1.s | p1.s | p1.s | p1.s |
| OPES MultiCV 1 | - | **D1**,yywo | **D1**,yywo | **D1**,yywo | **D1**,yywo | **D1**,yywo | **D1**,yywo | **D1**,yywo |
| OPES MultiCV 2 | - | - | **D2**,yywo | **D2**,yywo | **D2**,yywo | **D2**,yywo | **D2**,yywo | **D2**,yywo |
| OPES MultiCV 3 | - | - | - | **D3**,yywo | **D3**,yywo | **D3**,yywo | **D3**,yywo | **D3**,yywo |
| OPES MultiCV 4 | - | - | - | - | **D4**,yywo | **D4**,yywo | **D4**,yywo | **D4**,yywo |
| OPES MultiCV 5 | - | - | - | - | - | **D5**,yywo | **D5**,yywo | **D5**,yywo |
| OPES MultiCV 6 | - | - | - | - | - | - | **D6**,yywo | **D6**,yywo |
| OPES MultiCV 7 | - | - | - | - | - | - | - | **D7**,yywo |
| OPES MultiT | - | 301 K | 303 K | 306 K | 310 K | 317 K | 325 K | 335 K |

Table S1: The table presents the various CVs and parameters used in the *apo-MOR-OLD* OneOPES simulations. In the rows labelled "OPES Explore" and "OPES MultiCV", we outline the arrangement of the CVs across the replicas. The row titled "OPES MultiT" indicates the highest temperature reached during the trajectory, beginning from the thermostat's starting temperature of 300 K.

| Replicas | 0 | 1 | 2 | 3 | 4 | 5 | 6 | 7 |
|---|---|---|---|---|---|---|---|---|
| OPES Explore | Neop1.s | Neop1.s | Neop1.s | Neop1.s | Neop1.s | Neop1.s | Neop1.s | Neop1.s |
| OPES MultiCV 1 | - | **D1**,yywo | **D1**,yywo | **D1**,yywo | **D1**,yywo | **D1**,yywo | **D1**,yywo | **D1**,yywo |
| OPES MultiCV 2 | - | - | **D2**,yywo | **D2**,yywo | **D2**,yywo | **D2**,yywo | **D2**,yywo | **D2**,yywo |
| OPES MultiCV 3 | - | - | - | **D3**,yywo | **D3**,yywo | **D3**,yywo | **D3**,yywo | **D3**,yywo |
| OPES MultiCV 4 | - | - | - | - | **D4**,yywo | **D4**,yywo | **D4**,yywo | **D4**,yywo |
| OPES MultiCV 5 | - | - | - | - | - | **D5**,yywo | **D5**,yywo | **D5**,yywo |
| OPES MultiCV 6 | - | - | - | - | - | - | **D6**,yywo | **D6**,yywo |
| OPES MultiCV 7 | - | - | - | - | - | - | - | **D7**,yywo |
| OPES MultiT | - | 301 K | 303 K | 306 K | 310 K | 317 K | 325 K | 335 K |

Table S2: The table presents the various CVs and parameters used in the *apo-MOR-Euclidean* OneOPES simulations. In the rows labelled "OPES Explore" and "OPES MultiCV", we outline the arrangement of the CVs across the replicas. The row titled "OPES MultiT" indicates the highest temperature reached during the trajectory, beginning from the thermostat's starting temperature of 300 K.

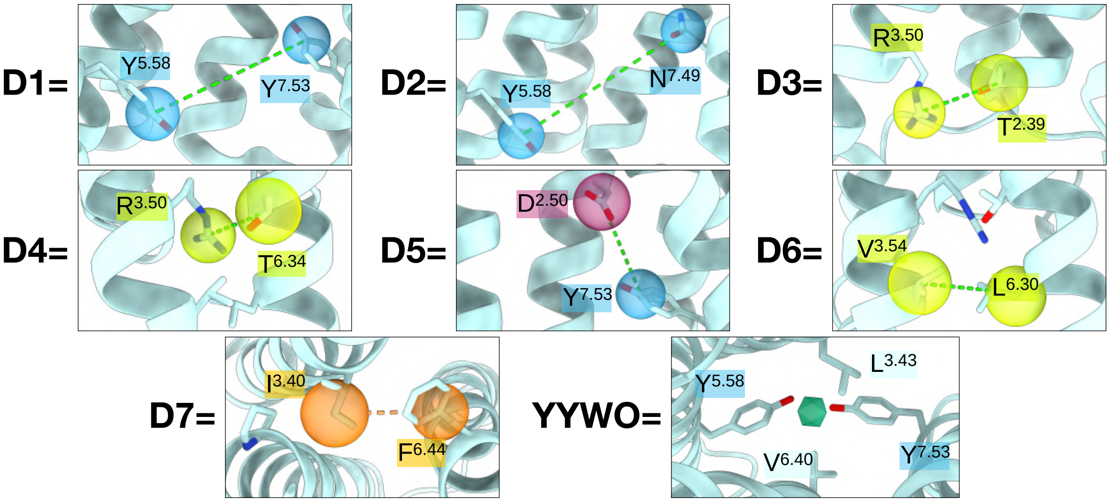


Figure S2: **Atomistic details of the CVs employed in the *apo-MOR-OLD* and *apo-MOR-Euclidean* OneOPES simulations.** For **D1-D7**, each CV is built as a distance between dummy atoms centered on the side-chains. The dummy atom built between $L^{3.43}$, $V^{6.40}$, and $Y^{7.53}$ and coordinating water molecules is represented as a green icosahedron.

**A)** ***Plumed.dat*:**

```
r3: RMSD REFERENCE=active_onlyCA.pdb TYPE=OPTIMAL
r4: RMSD REFERENCE=inactive_onlyCA.pdb TYPE=OPTIMAL
Neop1: PATH TYPE=EUCLIDEAN REFERENCE=MOR_euclidean_path.pdb LAMBDA=500.0
```

**B)** ***MOR_euclidean_path.pdb*:**

```
REMARK ARG=r4,r3 r4=0.1 r3=0.3
END
REMARK ARG=r4,r3 r4=0.15 r3=0.275
END
REMARK ARG=r4,r3 r4=0.18 r3=0.25
END
REMARK ARG=r4,r3 r4=0.225 r3=0.22
END
REMARK ARG=r4,r3 r4=0.25 r3=0.18
END
REMARK ARG=r4,r3 r4=0.28 r3=0.15
END
REMARK ARG=r4,r3 r4=0.3 r3=0.1
END
```

Figure S3: **How to setup the EPATH CV (upon which Neop1.s and Neop1.z are built) used in the MOR OneOPES simulations in this study. a)** Construction of the EPATH CV from the RMSD to the inactive (*r4*) and active (*r3*) reference structures, where milestones are defined exclusively through RMSD tuples rather than explicit intermediate conformations. **b)** Example of the reference file (with *.pdb* suffix) containing the tuple-defined milestones. This file must be provided to the PATH CV via the *REFERENCE=* keyword in the plumed.dat input.

### Supplementary Data 3

The following section displays both the construction of the sampled Euclidean PATH CV during the apo-ADRB1 (EPATH* hereafter), and the behaviour of the EPATH CV upon which the outcomes coming from *apo-ADRB1-Euclidean* have been re-weighted. In detail, we report here the full characterisation of the original CV definition. First, we show the set of milestone tuples in the ($RMSD_{\text{inactive}}$, $RMSD_{\text{active}}$) space used to build the EPATH* and EPATH (see Fig. S4A). We then provide the corresponding one-dimensional free-energy profile projected along the EPATH* CV, illustrating the relative stability of the inactive and active states (see Fig. S4B). Finally, we display the temporal evolution of the EPATH* value sampled by replica **0**, highlighting the extent of exploration and the convergence behaviour with the OneOPES protocol (see Fig. S4C).

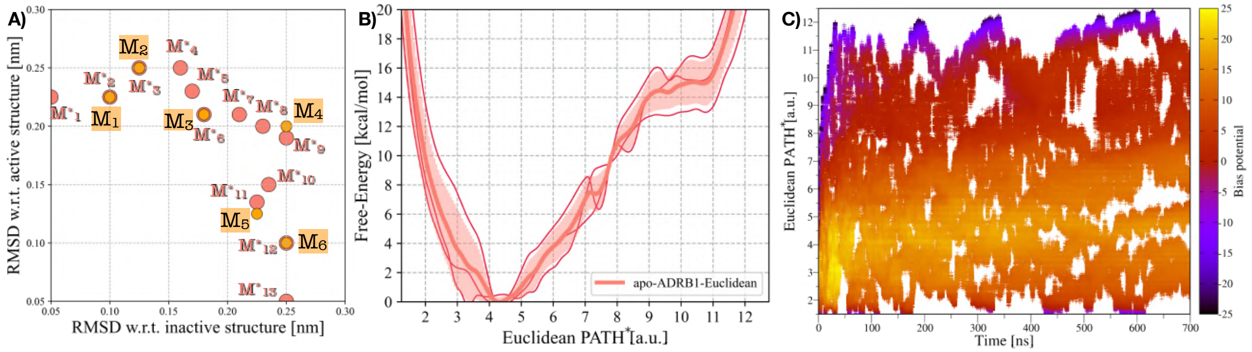


Figure S4: **Free-energy landscape and CV sampling in *apo-ADRB1-Euclidean* OneOPES simulations. a)** Milestones $M_i^*$ sampled during the *apo-ADRB1-Euclidean* OneOPES simulations (i.e. the sampled EPATH*), and milestones $M_i$ of the EPATH CV upon which the outcomes coming from *apo-ADRB1-Euclidean* have been re-weighted. **b)** Free-energy profile as a function of the apo-ADRB1 EPATH*, averaged over three independent OneOPES simulations. The solid pink line represents the mean free-energy, while the transparent pink shading indicates the standard deviation. **c)** Sampling of the EPATH* CV as a function of time in replica **0**, with data points colored according to the accumulated bias potential.

### Supplementary Data 4

In the following section, we report complementary analyses from the apo-ADRB1 Euclidean OneOPES simulations, focusing on the structural micro-switches that orchestrate GPCR activation. To elucidate how these local motifs couple to the global progression along the EPATH CV, we present a series of two-dimensional FES. These maps describe ADRB1's activation pathway in relation to the PIF, DRY, and YY micro-switches, as well as to the hydration state of the intracellular cavity. Together, these results provide a detailed view of how side-chain rearrangements and water-mediated interactions evolve in concert with the global transition from inactive to active conformations.

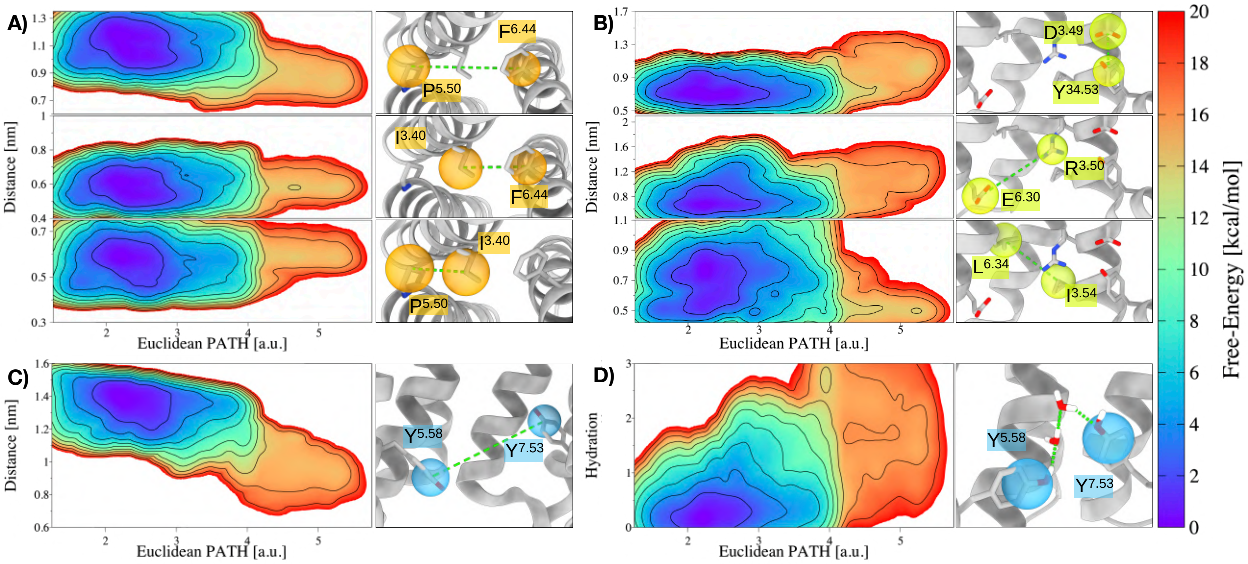


Figure S5: **Analyses of the micro-switches' behaviour in the *apo-ADRB1-Euclidean* OneOPES simulations**. **a)** 2D FES as a function of the EPATH CV and the PIF distances, illustrating the coupling between receptor progression and the PIF micro-switch. **b)** 2D FES as a function of the EPATH CV and the DRY distances, emphasising the structural transitions associated with the ionic lock $R^{3.50}$-$E^{6.30}$. **c)** 2D FES as a function of the EPATH CV and the YY distance, providing insights into the role of the YY micro-switch along the activation pathway. **d)** A fourth 2D FES illustrating the EPATH CV in relation to the hydration of the intracellular cavity.

### Supplementary Data 5

In this section, we provide additional data about the behaviour of the RPATH CV and the original EPATH* CV in the context of the apo-MOR OneOPES simulations. First, we report the temporal sampling of the RPATH CV, the RMSD PATH CV obtained by performing steered MD simulations starting from the inactive state of MOR (as in PDB ID: 9MQJ) and targeting the C$\alpha$s of MOR in an active conformation (PDB ID: 8F7R[14]). Fig. S6 illustrates how the traditional formulation explores the activation landscape in the *apo-MOR-OLD* setup. We then present the corresponding sampling of the EPATH* CV from the *apo-MOR-Euclidean* simulations, also including the set of Euclidean milestones $M_i^*$ used to construct the reference path in the ($RMSD_{inactive}$, $RMSD_{active}$) plane (see Fig. S7).

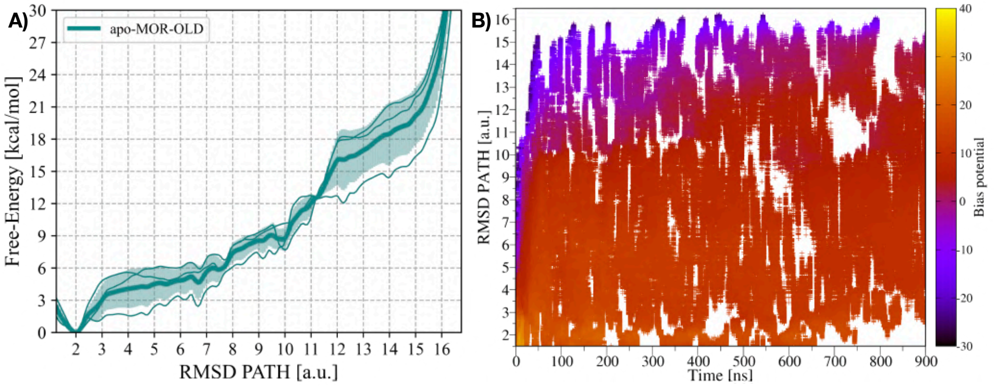


Figure S6: **Free-energy landscape and CV sampling in *apo-MOR-OLD* OneOPES simulations. a)** Free-energy profile as a function of the apo-MOR RPATH, averaged over three independent OneOPES simulations. The solid teal line represents the mean free-energy, while the transparent teal shading indicates the standard deviation. **b)** Sampling of the RPATH CV as a function of time in replica **0**, with data points colored according to the accumulated bias potential.

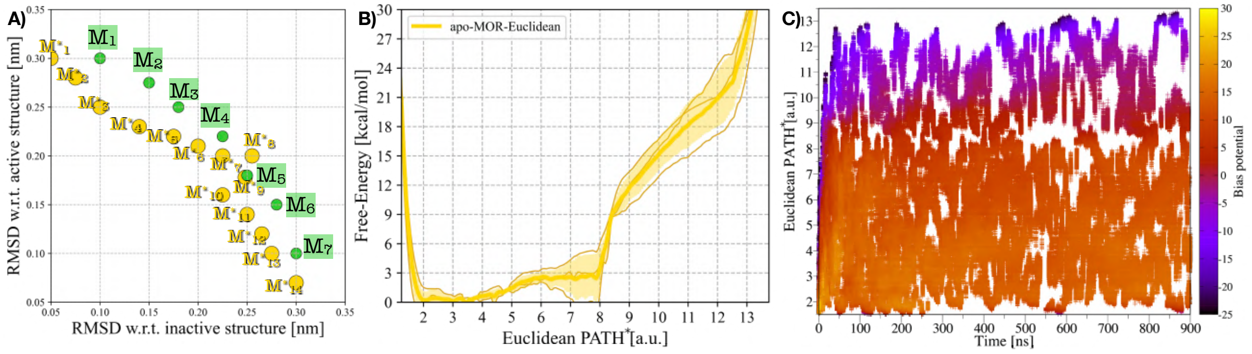


Figure S7: **Free-energy landscape and CV sampling in *apo-MOR-Euclidean* OneOPES simulations. a)** Milestones $M_i^*$ sampled during the *apo-MOR-Euclidean* OneOPES simulations (i.e. the sampled EPATH*), and milestones $M_i$ of the EPATH CV upon which the outcomes coming from *apo-MOR-Euclidean* have been re-weighted. **b)** Free-energy profile as a function of the apo-MOR EPATH*, averaged over three independent OneOPES simulations. The solid pink line represents the mean free-energy, while the transparent pink shading indicates the standard deviation. **c)** Sampling of the EPATH* CV as a function of time in replica **0**, with data points colored according to the accumulated bias potential.

#### Supplementary Data 6

In this section, we report a comparative analysis of MOR micro-switch rearrangements obtained from the *apo-MOR-OLD* and *apo-MOR-Euclidean* OneOPES simulations. For each microswitch (i.e., the *NPxxY*, the *DRY*, and the *YY* motifs), we computed 2D FES as a function of the receptor's activation coordinate EPATH and the corresponding structural descriptor (see Fig. S8). At the same time, we also monitored the behaviour of water molecules, whose accumulation in the intracellular cavity of the GPCR is a circumstance favouring the activation of the receptor (see Fig. S9). To deliver reliable statistics, the reported 2D FES are coming from the averages of the independent replicas. The collected results show a clear qualitative and quantitative agreement between *apo-MOR-OLD* and *apo-MOR-Euclidean*, further reinforcing that the new tuple-based approach reliably reproduces the hallmark micro-switch behaviour while offering a more robust and user-friendly construction of the activation path.

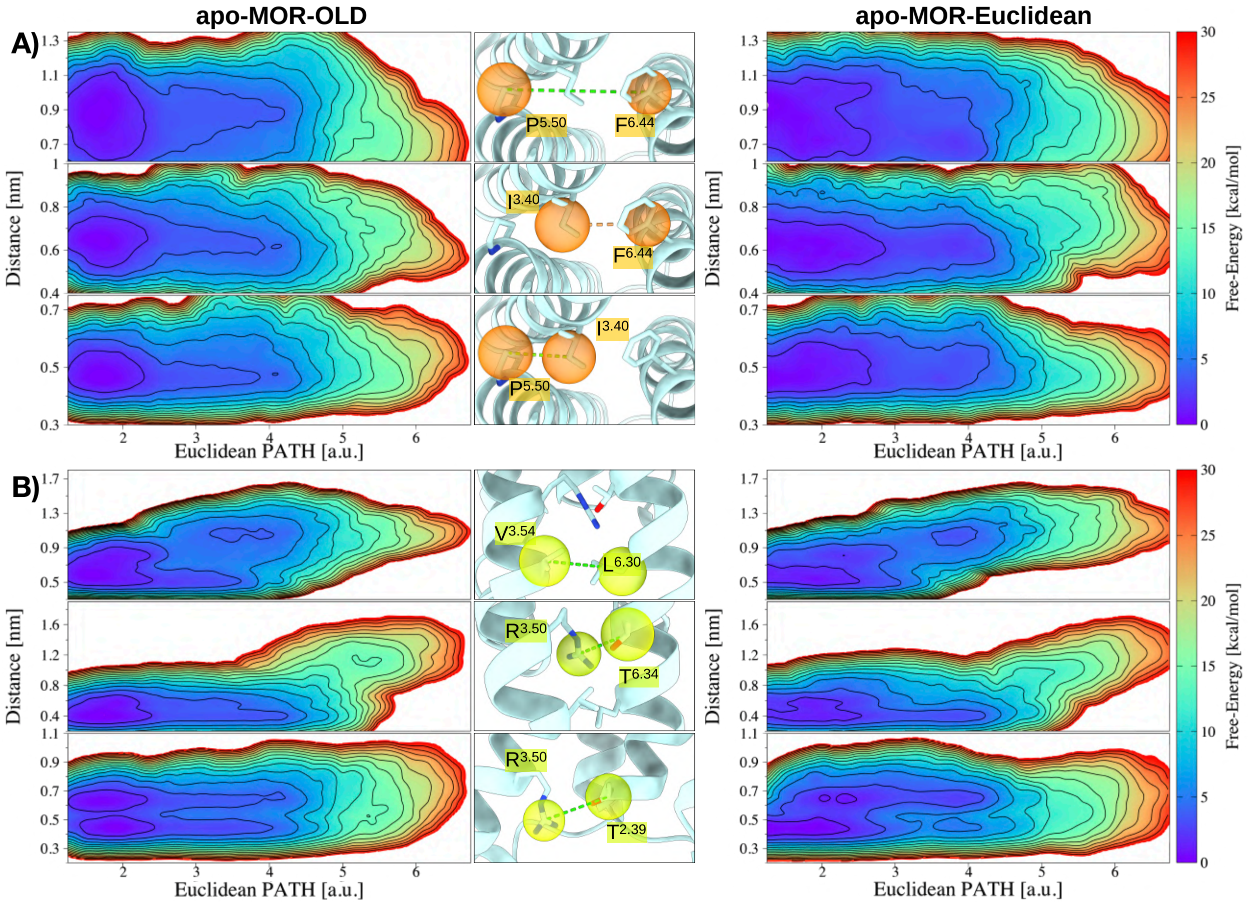


Figure S8: **Analyses of MOR's microswitches during the *apo-MOR-OLD* and *apo-MOR-Euclidean* OneOPES simulations. a)** 2D FES monitoring the Distance CVs for the residues of the *PIF* motif (i.e., $P^{5.50}$–$F^{6.44}$, $I^{3.40}$–$F^{6.44}$, and $P^{5.50}$–$I^{3.40}$) as a function of the EPATH CV for *apo-MOR-OLD* (left panel) and *apo-MOR-Euclidean* (right panel). **b)** 2D FES monitoring the Distance CVs for the residues of the *DRY* motif (i.e., $V^{2.54}$–$L^{6.30}$, $R^{3.50}$–$T^{6.34}$, and $R^{3.50}$–$T^{2.39}$) as a function of the EPATH CV (right panel) for *apo-MOR-OLD* (left panel) and *apo-MOR-Euclidean* (right panel). Isolines are drawn every 2 kcal/mol.

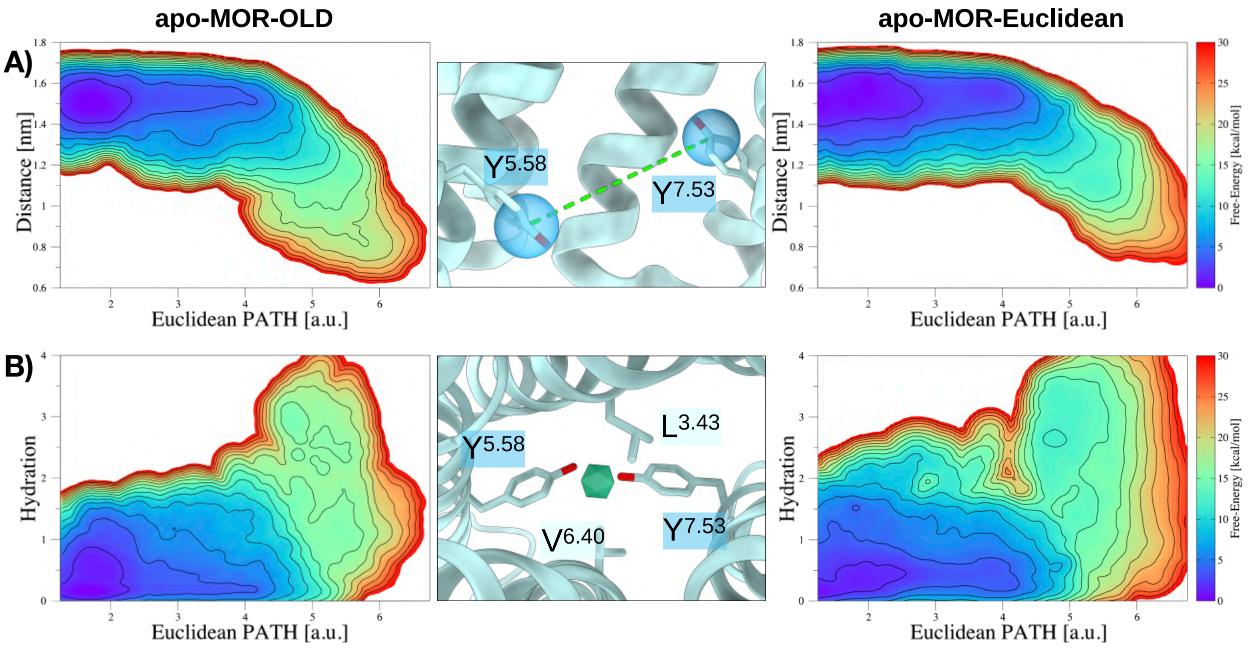


Figure S9: **Analysis of MOR's YY-motif and the hydration of the intracellular cavity. a)** 2D FES monitoring the Distance CVs for the residues of the *YY* motif (i.e., $Y^{5.58}$–$Y^{7.53}$) as a function of the EPATH CV for *apo-MOR-OLD* (left panel) and *apo-MOR-Euclidean* (right panel). **b)** 2D FES associated with the hydration of the *YY*-motif during the GPCR activation, as a function of the EPATH CV for *apo-MOR-OLD* (left panel) and *apo-MOR-Euclidean* (right panel). Isolines are drawn every 2 kcal/mol.

### Supplementary Data 7

In this section, we provide additional details on the characterisation of the conformational sampling achieved in *apo-MOR-OLD* and *apo-MOR-Euclidean* simulations as displayed in Fig. 4.

To visualise how the system explores the activation landscape, we project the trajectories onto the ($RMSD_{inactive}$; $RMSD_{active}$) plane and examine the distribution of frames associated with each Euclidean PATH interval (M1–M2 through M6–M7). This representation allows a direct comparison between the traditional *apo-MOR-OLD* and the tuple-based *apo-MOR-Euclidean*, highlighting their consistency in sampling the GPCR's progression from inactive to active–like states. In addition, we report histograms showing the frequency of occurrence of the clusters identified within each milestone interval.

The clustering analysis reveals that the terminal milestones (M1–M2 and M6–M7) are dominated by a single, highly populated cluster family, whereas the intermediate milestones exhibit multiple coexisting families, reflecting the increased conformational freedom and heterogeneity characteristic of the central region of the activation pathway.

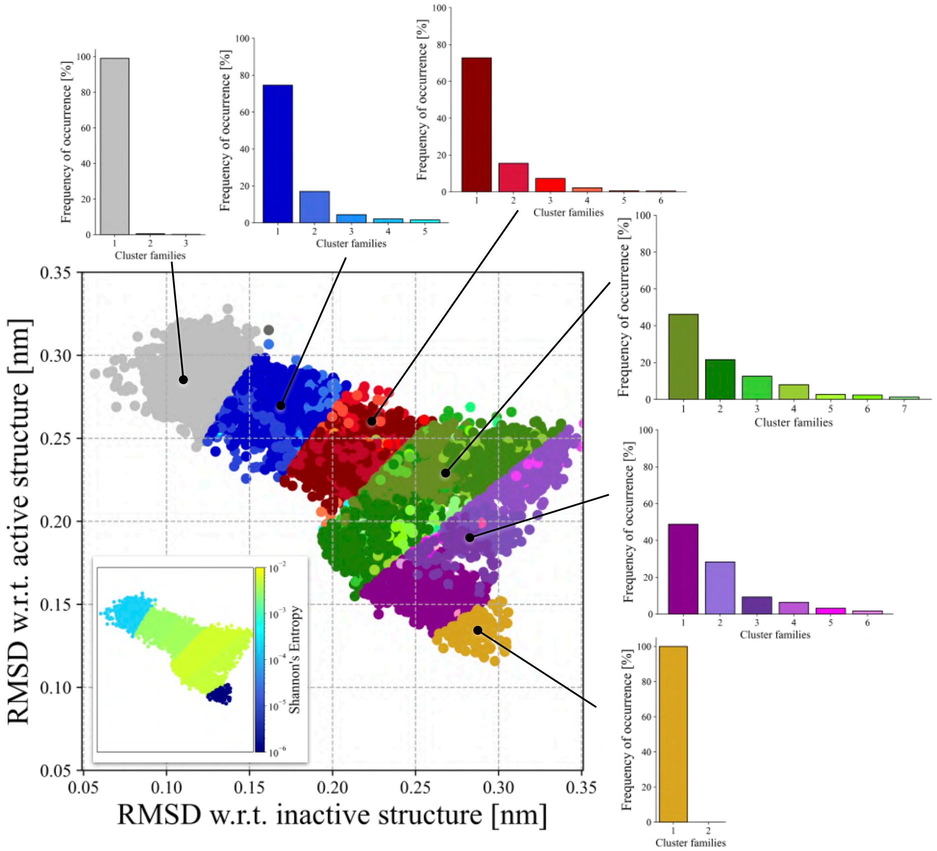


Figure S10: **Sampling along the EPATH CV in the *apo-MOR-OLD* simulation.** Sampling distribution projected onto the ($RMSD_{inactive}$; $RMSD_{active}$) space. Frames are grouped according to their EPATH intervals (M1–M2, M2–M3, M3–M4, M4–M5, M5–M6, M6–M7), with points in each interval coloured according to their assigned cluster (grey, blue, red, green, purple, and yellow palettes, respectively). The frequencies of occurrence of each cluster family are shown through histograms, coloured following the same colour palette. An inset inside the plot shows the same point distribution coloured by the local Shannon entropy.

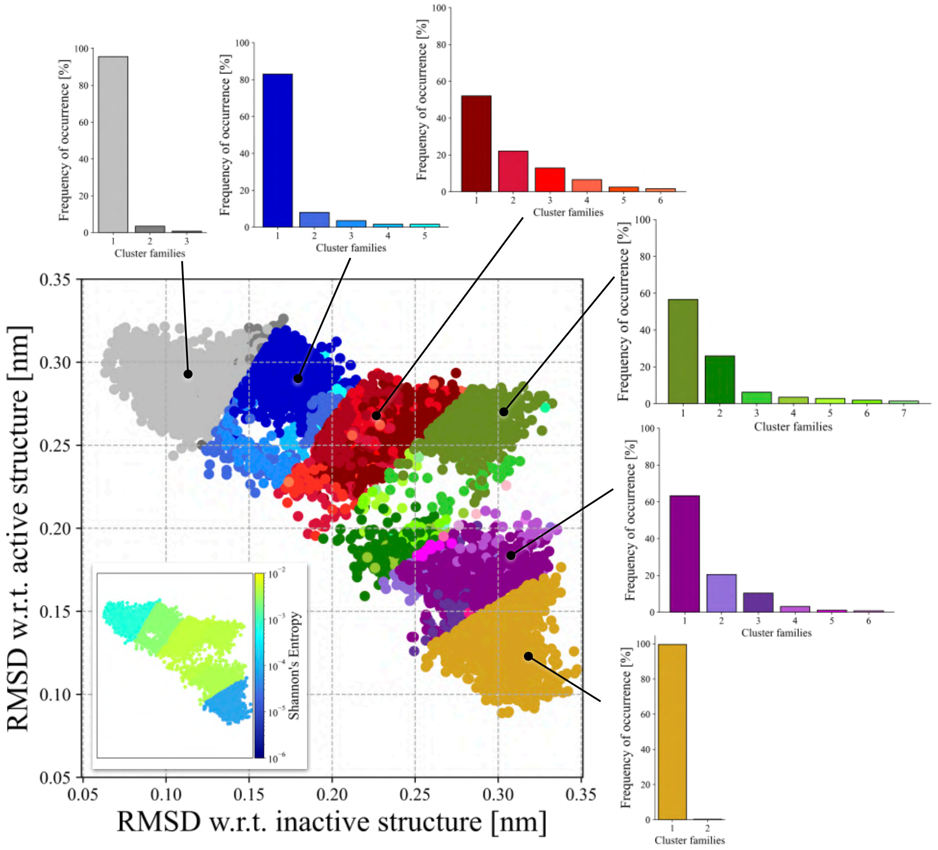


Figure S11: **Sampling along the EPATH CV in the *apo-MOR-Euclidean* simulation.** Sampling distribution projected onto the ($RMSD_{inactive}$; $RMSD_{active}$) space. Frames are grouped according to their EPATH intervals (M1–M2, M2–M3, M3–M4, M4–M5, M5–M6, M6–M7), with points in each interval coloured according to their assigned cluster (grey, blue, red, green, purple, and yellow palettes, respectively). The frequencies of occurrence of each cluster family are shown through histograms, coloured following the same colour palette. An inset inside the plot shows the same point distribution coloured by the local Shannon entropy.

#### References

[1] Seth F Vigneron, Shohei Ohno, Joao Braz, Joseph Y Kim, Oh Sang Kweon, Chase Webb, Christian B Billesbølle, Karthik Srinivasan, Karnika Bhardwaj, John J Irwin, et al. Docking 14 million virtual isoquinuclidines against the $\mu$ and $\kappa$ opioid receptors reveals dual antagonists–inverse agonists with reduced withdrawal effects. *ACS Central Science*, 11(5):770–790, 2025.

[2] Elaine C Meng, Thomas D Goddard, Eric F Pettersen, Greg S Couch, Zach J Pearson, John H Morris, and Thomas E Ferrin. Ucsf chimerax: Tools for structure building and analysis. *Protein Science*, 32(11):e4792, 2023.

[3] Eric F Pettersen, Thomas D Goddard, Conrad C Huang, Elaine C Meng, Gregory S Couch, Tristan I Croll, John H Morris, and Thomas E Ferrin. Ucsf chimerax: Structure visualization for researchers, educators, and developers. *Protein science*, 30(1):70–82, 2021.

[4] Sunhwan Jo, Xi Cheng, Jumin Lee, Seonghoon Kim, Sang-Jun Park, Dhilon S Patel, Andrew H Beaven, Kyu Il Lee, Huan Rui, Soohyung Park, et al. Charmm-gui 10 years for biomolecular modeling and simulation. *Journal of computational chemistry*, 38(15):1114–1124, 2017.

[5] Stefano Piana, Paul Robustelli, Dazhi Tan, Songela Chen, and David E. Shaw. Development of a force field for the simulation of single-chain proteins and protein-protein complexes. *Journal of Chemical Theory and Computation*, 16:2494–2507, 2020.

[6] Mark James Abraham, Teemu Murtola, Roland Schulz, Szilárd Páll, Jeremy C Smith, Berk Hess, and Erik Lindahl. Gromacs: High performance molecular simulations through multi-level parallelism from laptops to supercomputers. *SoftwareX*, 1:19–25, 2015.

[7] Berk Hess, Henk Bekker, Herman JC Berendsen, and Johannes GEM Fraaije. Lincs: A linear constraint solver for molecular simulations. *Journal of computational chemistry*, 18(12):1463–1472, 1997.

[8] Berk Hess. P-lincs: A parallel linear constraint solver for molecular simulation. *Journal of chemical theory and computation*, 4(1):116–122, 2008.

[9] Henrik G Petersen. Accuracy and efficiency of the particle mesh ewald method. *The Journal of chemical physics*, 103(9):3668–3679, 1995.

[10] Giovanni Bussi, Davide Donadio, and Michele Parrinello. Canonical sampling through velocity rescaling. *The Journal of chemical physics*, 126(1):014101, 2007.

[11] Mattia Bernetti and Giovanni Bussi. Pressure control using stochastic cell rescaling. *The Journal of Chemical Physics*, 153(11):114107, 2020.

[12] Simone Aureli, Valerio Rizzi, Nicola Piasentin, and Francesco Luigi Gervasio. Enhanced sampling and tailored collective variables yield reproducible free energy landscapes of beta-1 adrenergic receptor activation. *Journal of Chemical Theory and Computation*, 21(15):7687–7700, 2025.

[13] Xinyu Xu, Jonas Kaindl, Mary J Clark, Harald Hübner, Kunio Hirata, Roger K Sunahara, Peter Gmeiner, Brian K Kobilka, and Xiangyu Liu. Binding pathway determines norepinephrine selectivity for the human $\beta$1ar over $\beta$2ar. *Cell Research*, 31(5):569–579, 2021.

[14] Yue Wang, Youwen Zhuang, Jeffrey F DiBerto, X Edward Zhou, Gavin P Schmitz, Qingning Yuan, Manish K Jain, Weiyi Liu, Karsten Melcher, Yi Jiang, et al. Structures of the entire human opioid receptor family. *Cell*, 186(2):413–427, 2023.
